# Unique and overlapping behavioral effects of isoform-specific NRXN1 deletions across development

**DOI:** 10.64898/2026.01.22.698702

**Authors:** Amanda E. Friedman, Michele Perni, Josephine Millard, Damjan Karanfilovski, Michael Granato, Philip D. Campbell

## Abstract

Mutations in the human neurexin 1 (NRXN1) gene are associated with neurodevelopmental disorders including autism and schizophrenia. Vertebrate NRXN1 produces three major NRXN1 isoforms, named *α, β*, and *γ*. Human genetic data suggests that deletions located at the 5’ region of the gene disrupting the *α* isoform associate mostly with clinical behavioral deficits. Yet, 3’ deletions that disrupt multiple isoforms have also been identified in clinical cases. Whether deletions that selectively affect specific NRXN1 isoforms result in specific behavioral deficits remains unclear. Here, we show that zebrafish harboring gene deletions that differentially encompass *nrxn1a-α, -β*, and/or *-γ* display both unique and overlapping locomotion, sensorimotor behavior, and social behavior deficits. Combined, our results strongly support a model by which domain selective *nrxn1* deletions predict behavioral phenotypes. Moreover, our results demonstrate compelling relationships between genotype and resulting behavioral phenotypes, providing functional insights into the complexity of NRXN1-associated neuropsychiatric behaviors.

## Introduction

Neurexin-1 (NRXN1) is a pre-synaptic adhesion molecule that plays important roles in synapse formation, maintenance, regulation, and function^1^. The *NRXN1* locus is particularly susceptible to non-recurrent copy-number variants (i.e. unique genomic changes that occur at different locations in different individuals), and *NRXN1* deletions have been associated with multiple neurodevelopmental disorders including autism, schizophrenia, intellectual disability, and developmental delay^2^. While most individuals ascertained to date have monoallelic deletions, biallelic *NRXN1* deletions have been observed in a small group of individuals diagnosed with Pitt-Hopkins syndrome who display significant development delay^2,3^.

Genomic studies have described three different promoters at the human and mouse NRXN1 loci that generate three evolutionary conserved *NRXN1* isoforms known as *α, β*, and *γ*^*1*^. While their intracellular domains are identical, their extracellular domains differ, with the *α* isoform being the longest, the *β* isoform being intermediate in length and the *γ* isoform being the shortest. To date, *α*-specific exonic deletions located in the 5’ region of NRXN1 have been strongly associated with clinically defined behavioral phenotypes^4^, suggesting that, compared to β or γ deletions, disruption of the *NRXN1-α* isoform may have a major impact on behavior and thus be more pathogenic. However, deletions involving the β isoform have also been identified in individuals with clinical phenotypes^2,4^, suggesting a more nuanced view. Indeed, recent work has suggested that 5’ and 3’ *NRXN1* deletions can act through divergent molecular mechanisms^5,6^ requiring altogether different interventions. Therefore, understanding if and to which degree NRXN1 isoform specific deletions impact behavioral outcomes is essential to define the spectrum of disease mechanisms. This knowledge in turn may ultimately predict more fine-tuned and divergent therapeutic interventions.

To date, most behavioral studies of NRXN1 isoforms have focused on mammalian *NRXN1-α*. These studies reveal that in mice, homozygous *NRXN1-α*-specific deletions cause multiple behavioral abnormalities including alterations in social behavior, locomotion, and anxiety^7–9^ that are inconsistently seen across studies in heterozygous animals^7,8,10,11^.

Surprisingly, analogous studies evaluating the function of the NRXN1 *β* or *γ* isoforms and whole-gene NRXN1 deletions spanning all the isoforms have been missing.

To fill this gap, we used zebrafish, which unlike *Drosophila* and *C. elegans* has maintained a mammalian-like tripartite *nrxn1* promoter architecture. Here, leveraging the strengths of the zebrafish system, we generate zebrafish *nrxn1* whole gene and isoform-specific deletion lines and assess how these deletions affect sensorimotor and social behavior across development. Our results demonstrate that isoform-specific *nrxn1* deletions result in both shared and unique deficits that are both behaviorally and temporally specific. Early in development, we identify opposing sensorimotor phenotypes caused by *nrxn1-α*-specific and *nrxn1-β*-specific deletions. Later in development, we show that *nrxn1* is required for social behavior but show that *nrxn1-α*-specific deletions do not fully recapitulate *nrxn1* deletions spanning all isoforms. Together, our data reveal that NRXN1 deletions that differentially encompass *α, β*, and *γ* produce different behavioral outcomes, supporting a more complex and nuanced view of how distinct *nrxn1* mutations predict selective neurodevelopmental phenotypes.

## Results

### *nrxn1a* crispants display robust sensorimotor behavioral phenotypes

The zebrafish genome possesses two copies of *NRXN1*, encoded by two genetically distinct loci, *nrxn1a* on chromosome 12 and *nrxn1b* on chromosome 13, respectively (Fig 1A,B). At the protein level, both genes are highly similar to human NRXN1 (similarity/identify 76%/85% and 68%/78%, respectively)^12^ and are predicted to have identical protein architectures compared to their mammalian counterparts (Fig 4A,B). To determine the neurodevelopmental function of each gene, we selectively disrupted *nrxn1a* or *nrxn1b* using a CRISPR-Cas9-based approach that has been previously reported to lead to biallelic null alleles in over 90% of guide RNA injected animals^13^. Briefly, three guide RNAs that target three non-overlapping sites along the *nrxn1a* and *nrxn1b* genes (Fig 1A,B) were injected into fertilized wild-type embryos together with Cas9 protein to generate *nrxn1a* and *nrxn1b* crispants. Embryos that were injected with three non-targeting gRNAs and Cas9 we used as controls. *nrxn1a* and *nrxn1b* crispants were viable at 6 days post-fertilization (dpf) and did not display obvious gross morphological defects (*nrxn1a*: n=117 crispants, n=169 controls; *nrxn1b*: n=148 crispants, n=124 controls; 3 biological replicates each). We next assessed motor behaviors of *nrxn1a* and *nrxn1b* crispants at 6dpf using our previously established and validated pipeline^14,15^ that allows assessment of multiple sensorimotor behaviors including the visual motor response (VMR)^16^, responsiveness to flashes of light (LF) or darkness (DF)^17,18^, and modulation of the acoustic startle response (ASR)^18–21^. Using this pipeline we failed to detect deficits for any of the behaviors assayed in *nrxn1b* crispants (Fig 1C. We later confirmed these results in *nrxn1b* whole gene deletion (*nrxn1b-WD*) mutants (Fig 4B, 5E,E’,J,O). In contrast, *nrxn1a* crispants displayed clear phenotypes that spanned multiple behaviors (Fig 1C). Specifically, *nrxn1a* crispants displayed reduced movement metrics to visual stimuli during the VMR (i.e. distance travelled, average speed, and average speed per bout) and DF (i.e. average distance and duration travelled following dark flash) assays, and increased sensitivity to acoustic stimuli in the ASR assay (i.e. percent short latency C-bend in response to loud acoustic stimuli and area under the curve (AUC)) (Fig 1C). Together, these results suggest that compared to *nrxn1b, nrxn1a* may play more dominant neurodevelopmental roles. Thus, for our isoform-specific analysis, we focused on the *nrxn1a* gene.

**Figure 1.**
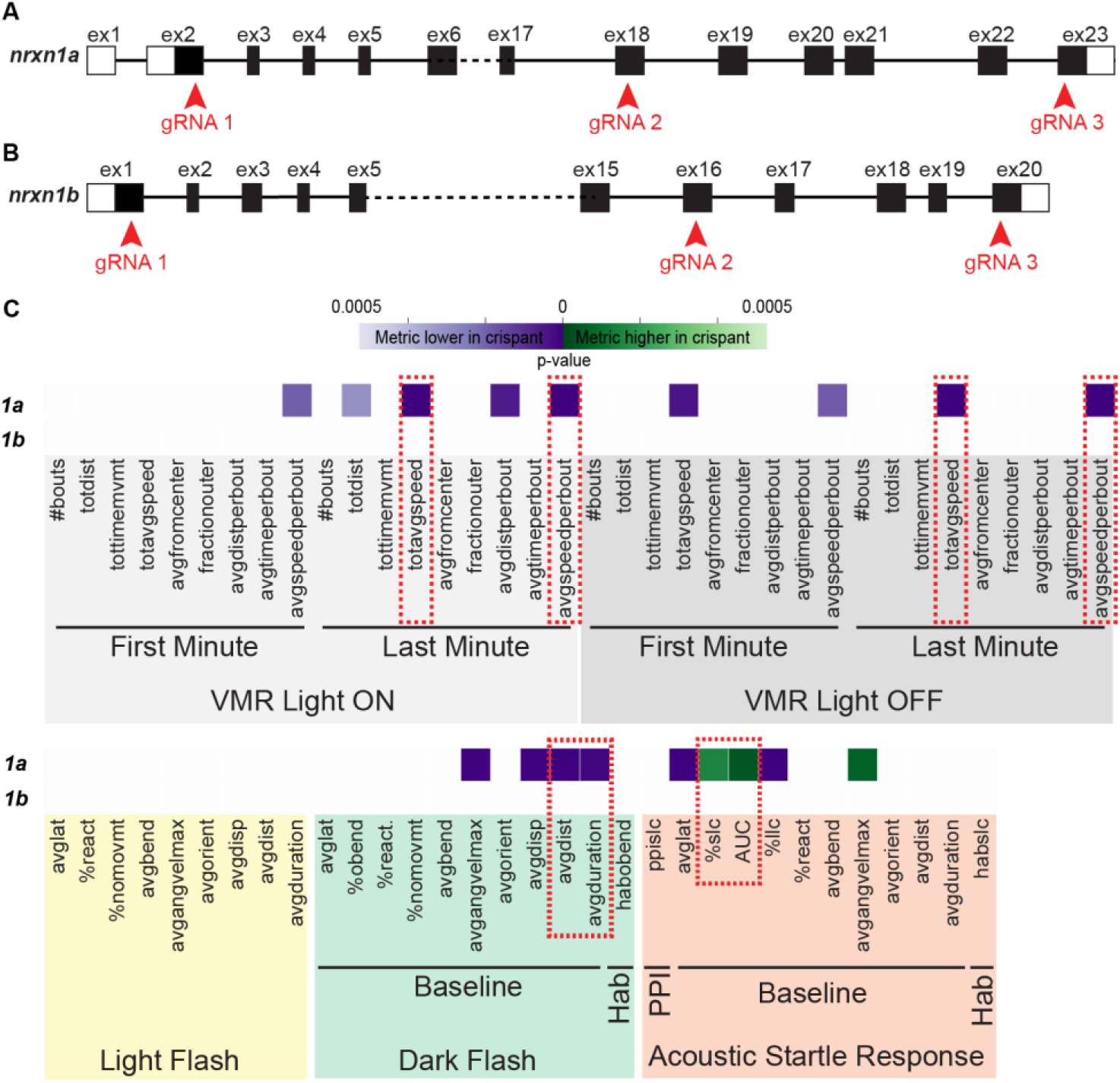
*nrxn1a* crispants display robust sensorimotor behavioral phenotypes. (A,B) Exon structure of nrxn1a (A) and nrxn1b (B) showing locations of gRNA targets for crispant generation. (C) Heatmap illustrating *nrxn1a* and *nrxn1b* crispant phenotypes across behavioral metrics. Each row is an individual crispant and all columns represent a behavioral metric. Boxes that are colored designate statistically significant difference in the metric in crispant vs controls based on Student’s t-test with a Bonferroni-corrected p-value of <0.05/94. VMR behavioral metrics are highlighted in gray, LF in yellow, DF in green, and ASR in orange. Red broken boxes denote reduced movement metrics during the VMR and DF and increased ASR sensitivity. For both genes, three biological replicates were performed. A total of n= 117 crispants and n=169 controls were assayed for *nrxn1a*. A total of n= 148 crispants and n=124 controls were assayed for *nrxn1b*. Presented data is representative from a single biological replicate.

### *nrxn1a-α* and *-β* have distinct brain expression patterns

In humans, NRXN1 has three main promoter-driven isoforms which are termed *α, β*, and *γ*. Consistent with the tripartite promoter architecture described in mammals, the zebrafish *nrxn1a* locus has retained three major H3K4me3 promoter peaks in ZebrafishENCODE brain data (Fig 2A)^22^. While Ensembl^23^ currently annotates only transcripts corresponding to the *α* and *β* isoforms, a third epigenomic peak supports the presence of an *γ*-like promoter. Indeed, Ensembl includes an additional RNA-seq–supported *nrxn1a* transcript in 5dpf larvae originating from a third promoter region, consistent with the *γ* promoter architecture (Fig 2B). Together, in accordance with the mammalian literature, this provides compelling evidence for three major promoter-driven *nrxn1a* isoforms which we similarly named *α, β*, and *γ* (Fig 2B).

**Figure 2.**
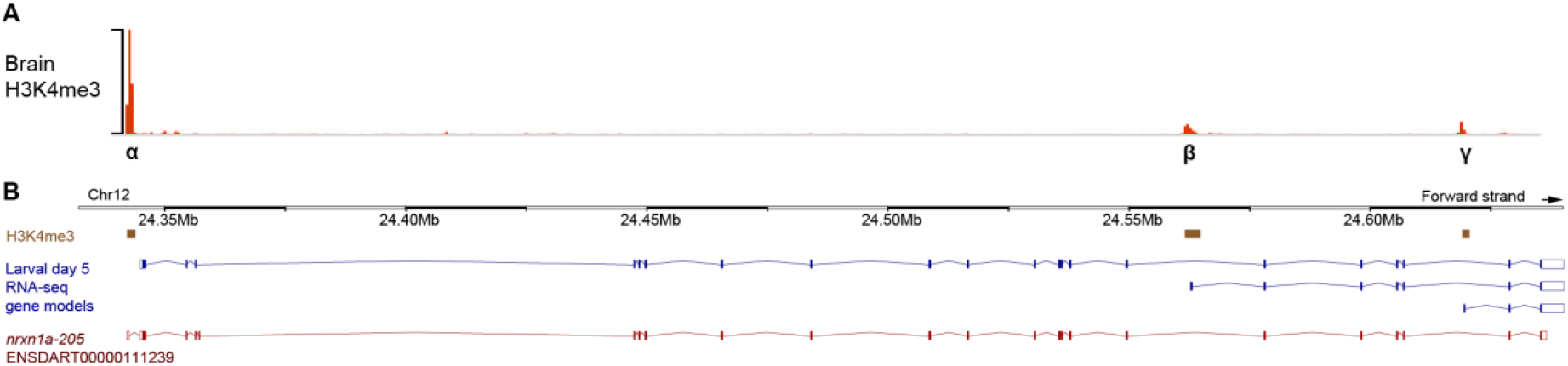
*nrxn1a* has three main promoter-defined transcripts. Transcript models from Ensembl (danRer11), supported by larval (5dpf) RNA-seq data (B), are shown relative to brain H3K4me3 enrichment from zebrafishENCODE (A).

To further validate the existence of the annotated *nrxn1a-α* and *β* isoforms and to define their expression patterns, we performed HCR in situ hybridization with probes specific for the *α* and *β* isoforms on brains at 6dpf to match our behavioral assessments. Both the *nrxn1a-α* and - *β* isoforms are widely expressed through the brain and display largely overlapping expression patterns (Fig 3). Despite the overall similarities, we detected an enrichment of *nrxn1a-β* mRNA signal in an anterior region of the forebrain bilaterally, corresponding to the pallium. *nrxn1a-α* also appeared to have a notably higher signal than *nrxn1a-β*, which correlates with the larger H3K4me3 peak and known higher expression of the *α* isoform in mammals^24^. Thus, in situ expression patterns could be consistent with a scenario by which *nrxn1a-α* and -*β* isoforms have both overlapping and unique neurodevelopmental functions that might result in behavioral diversity.

**Figure 3.**
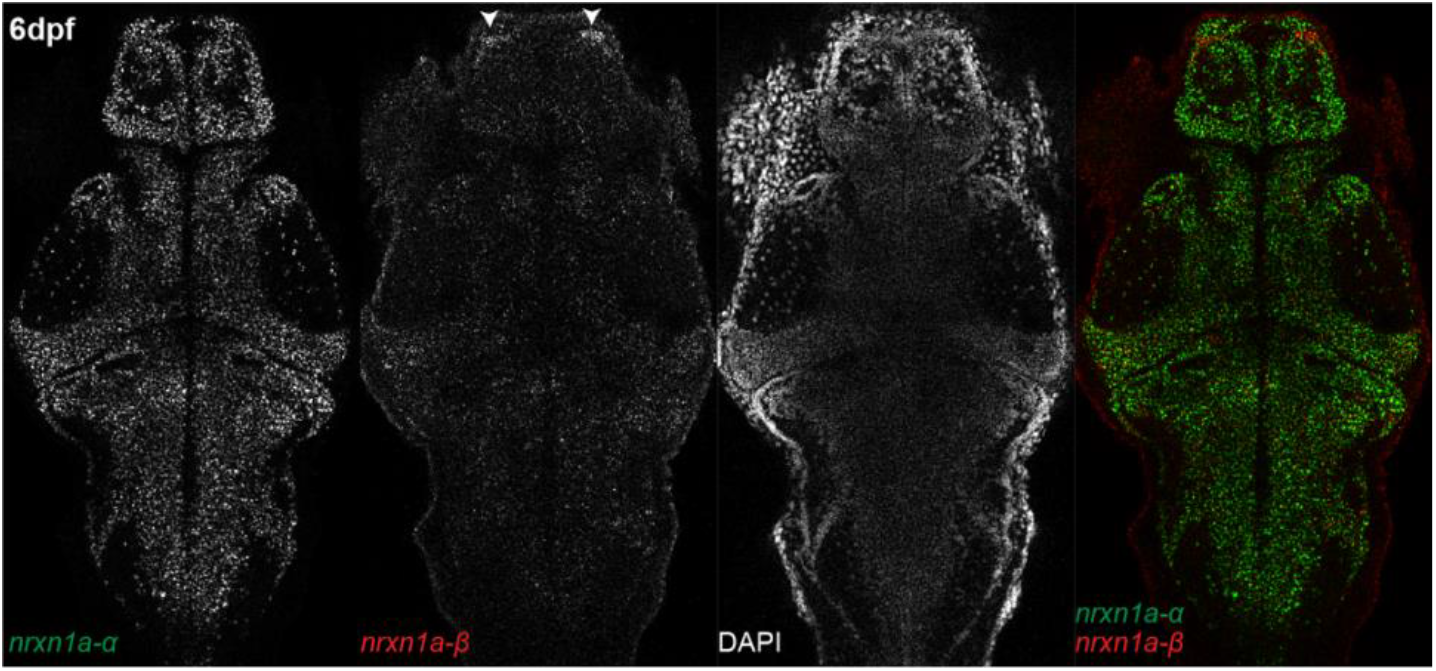
*nrxn1a-α* and *-β* have distinct brain expression patterns. HCR in situ hybridization of *nrxn1a* isoform-specific exons in wild-type larval brain at 6dpf. *nrxn1a-α* and *-β* largely overlap in expression, though *nrxn1a-β* shows enrichment in the pallium (arrowheads). Images are representative of n=12.

### Isoform-specific *nrxn1a* deletions cause distinct sensorimotor behavioral phenotypes

To define if and to which degree isoform-specific *NRXN1* deletions correlated with specific behavioral phenotypes, we used CRISPR-Cas9 mediated genome editing to generate isoform-specific *nrxn1a* deletion mutants. Specifically, we generated mutant lines harboring deletions that delete nearly the entire *nrxn1a* locus (*nrxn1a-WD*), and deletions spanning isoform-specific exons (*nrxn1a-α, nrxn1a-β, nrxn1a-γ*), respectively (Fig 4A). All the isoform-specific deletions are predicted to remove the start codon and to abolish the entire *nrxn1a* open reading frame, whereas the *nrxn1a-WD* deletion is predicted to delete all but a portion of the first LNS domain. As such, all of the deletions are predicted to cause loss of function.

**Figure 4.**
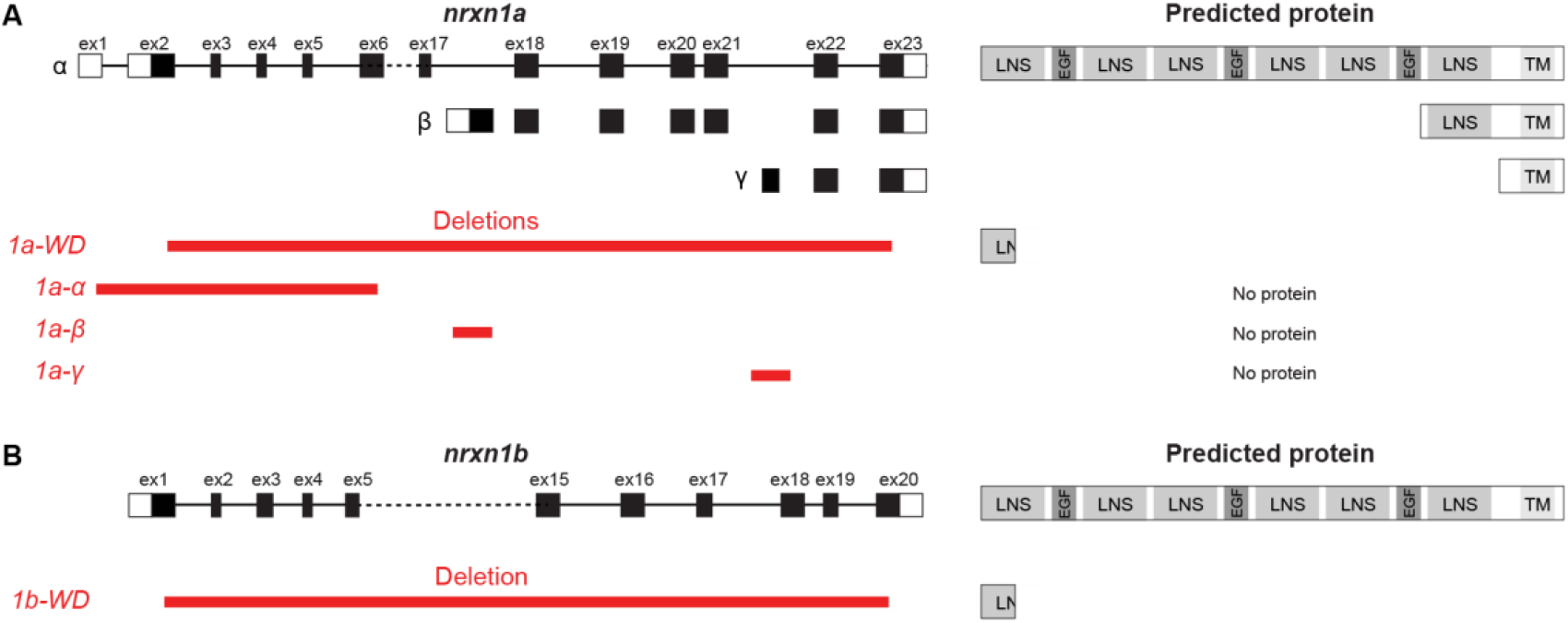
Zebrafish *nrxn1* deletion lines. (A) Exon structure of *nrxn1a* alongside *nrxn1a* deletion lines. *nrxn1a-WD* deletes nearly the entire locus, *nrxn1a-α*-specific line deletes *α*-specific exon, *nrxn1a-β*-specific line deletes the lone *β*-specific exon, and the *nrxn1a-γ*-specific line deletes the lone *γ*-specific exon. *nrxn1a-WD* is predicted to lead to a severely truncated protein with disruption of all major domains. *nrxn1a-α, -β*, and *-γ* lines are predicted to lead to no protein product. (B) Exon structure of *nrxn1b* alongside *nrxn1b* deletion line. *nrxn1b-WD* deletes nearly the entire locus and is predicted to lead to a severely truncated protein with disruption of all major domains.

For each mutant line, we then in-crossed heterozygous adults and analyzed behavior of wild-type, heterozygous, and homozygous offspring at 6dpf. Because *nrxn1a* crispants displayed reduced locomotion during the light-induced VMR and DF assays, and hypersensitivity of the ASR, we focused subsequent behavioral analyses on these behaviors. In all lines assayed, we did not observe differences between wildtypes and heterozygotes (Fig S1). Therefore, for our analysis, we compared homozygous mutants (*nrxn1a*^-/-^) to their combined homozygous wild-type (*nrxn1a*^+/+^) and heterozygous (*nrxn1a*^+/-^) siblings. Compared to siblings, *nrxn1a-WD* and *nrxn1a*-α mutants displayed reduced movement during the VMR assay, as defined by number of bouts, that was most pronounced during the light phase of the assay (first 8 minutes, Fig 5A,B), recapitulating the *nrxn1a* crispant results. We noted that, compared to *nrxn1a-WD*, the phenotype was more pronounced in *nrxn1a-α* mutants, which displayed a significant reduction in the number of bouts when averaging across the entire light period (Fig S2A,B). In contrast, both *nrxn1a-β* and *nrxn1a-γ* mutants displayed increased movement during the light phase of the VMR (Fig 5C,D). This phenotype was most pronounced in *nrxn1a-β* mutants which displayed a significant increase in the number of bouts when averaging across the entire light period (Fig S2C,D). This could explain why the *nrxn1a-WD* deletion phenotype, which also deletes *β* and *γ*, is less severe than the *nrxn1a-α* deletion phenotype. In the DF assay, *nrxn1a-WD* and *nrxn1a-α* mutants displayed reduced movement, defined by average distance travelled following a flash of darkness, compared to siblings (Fig 5F,G). In contrast, compared to siblings following a flash of darkness, *nrxn1a-β* mutants displayed a greater distance travelled (Fig 5H), while we failed to detect differences in *nrxn1a-γ* mutants (Fig 5I). In the ASR assay, compared to siblings, *nrxn1a-WD* mutants displayed a significant increase in sensitivity to startling stimuli (Fig 5K). No differences in the ASR were observed in *nrxn1a-α, nrxn1a-β*, or *nrxn1a-γ* mutants (Fig 5L-N), suggesting that while *nrxn1a* is critical to regulate ASR stimulus sensitivity, individual isoforms may functionally compensate for each other. Together, our results indicate that *in vivo* isoform-specific *nrxn1a* deletions cause distinct, and at times opposing, behavioral phenotypes. Further, we find that while *nrxn1a-α* deletion mimics the *nrxn1a-WD* VMR and DF phenotypes, the *nrxn1a-α* deletion fails to recapitulate the *nrxn1a-WD* ASR phenotype.

**Figure 5.**
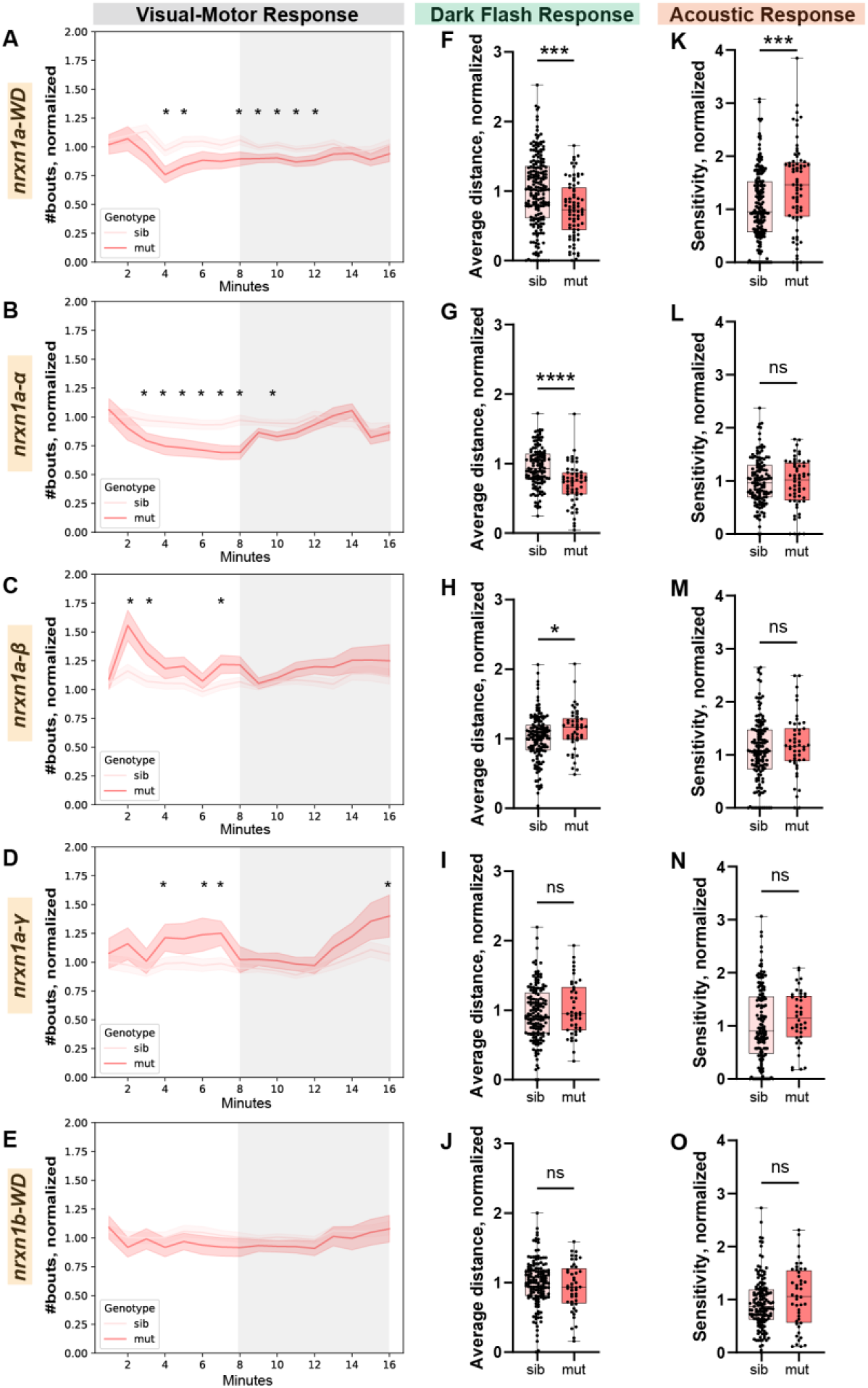
Isoform-specific *nrxn1a* deletions cause distinct sensorimotor behavioral phenotypes. (A-E) Number of bouts moved during each minute of the VMR assay, normalized to the wild-type average for each deletion line. White (first 8 minutes) and Gray (last 8 minutes) shading denote the light and dark periods of the VMR, respectively. The homozygous mutant (dark pink) and sibling (light pink) means +/-SEMs are plotted for each minute bin. Minutes where p<0.05 based on Student’s t-test are denoted with an asterisk (*). (A’-E’) Average of normalized bouts across all light (white) and dark (gray) minutes for each deletion line. (F-J) Average distance travelled following a flash of darkness, normalized to wildtypes for each deletion line. (K-O) Sensitivity to acoustic stimuli, normalized to wildtypes for each deletion line. *nrxn1a-WD*: n=75 mut, 197 sib (131 het, 66 wt); *nrxn1a-α*: n=56 mut, 130 sib (79 het, 51 wt); *nrxn1a-β*: n=47 mut, 149 sib (104 het, 45 wt); *nrxn1a-γ*: n=44 mut, 154 sib (96 het, 58 wt); *nrxn1b-WD*: n=41 mut, 143 sib (99 hets, 44 wt). Data is presented as cumulative across 4-5 biological replicates. *p<0.05, ***p<0.001, ****p<0.0001 based on Student’s t-test.

### nrxn1a deletions cause social interaction deficits

Given NRXN1’s strong association with human neurodevelopmental disorders such as autism, we next assayed social behaviors. For this, we developed an observer-independent, hands-free assay for the analysis of 5 weeks post-fertilization (wpf) juvenile fish that enables simultaneous imaging of baseline locomotion and social preference (Fig 6A-C). Briefly, we designed an arena composed of a long individual lane flanked on each end by a smaller compartment into which social stimulus fish or objects can be placed. Unlike most previously reported arenas^25–28^, our design visually separates the testing lane from the end compartments using microcontrolled polymer dispersed liquid crystal (PDLC) films (Fig 6C), of which opacity is controlled without any mechanical or moving parts that can interfere with behavior. To increase experimental throughput, we constructed a multiplex arena that consists of 14 individually controllable arenas in a 7×2 side-by-side configuration, allowing us to test 14 juvenile zebrafish at 5wpf robustly for social interaction activity (Fig 6A-C). For all experiments, we placed a ‘tester’ fish in the larger compartment of the arena, an age-matched ‘stimulus’ fish in one of the small compartments, and a novel object (black acrylic rectangular prism) in the other small compartment. Following placement into the testing arena and acclimation, movement and location of the tester fish are recorded with the PDLC windows switched off, or opaque, for 10mins, a period defined as the ‘baseline period’. Then, the PDLC windows are switched on, or transparent (open in Fig. 6), and fish are recorded for an additional 10mins as they respond to social and novel object stimuli (Fig 5D). During this ‘social period’, social preference (SP) is calculated for each fish by subtracting the fraction of time in the social zone during the baseline period from the fraction of time in the social zone during the social period. This provides a metric that falls within -1 and +1 with positive values indicating more time spent in the social zone during the social period than during the baseline period.

**Figure 6.**
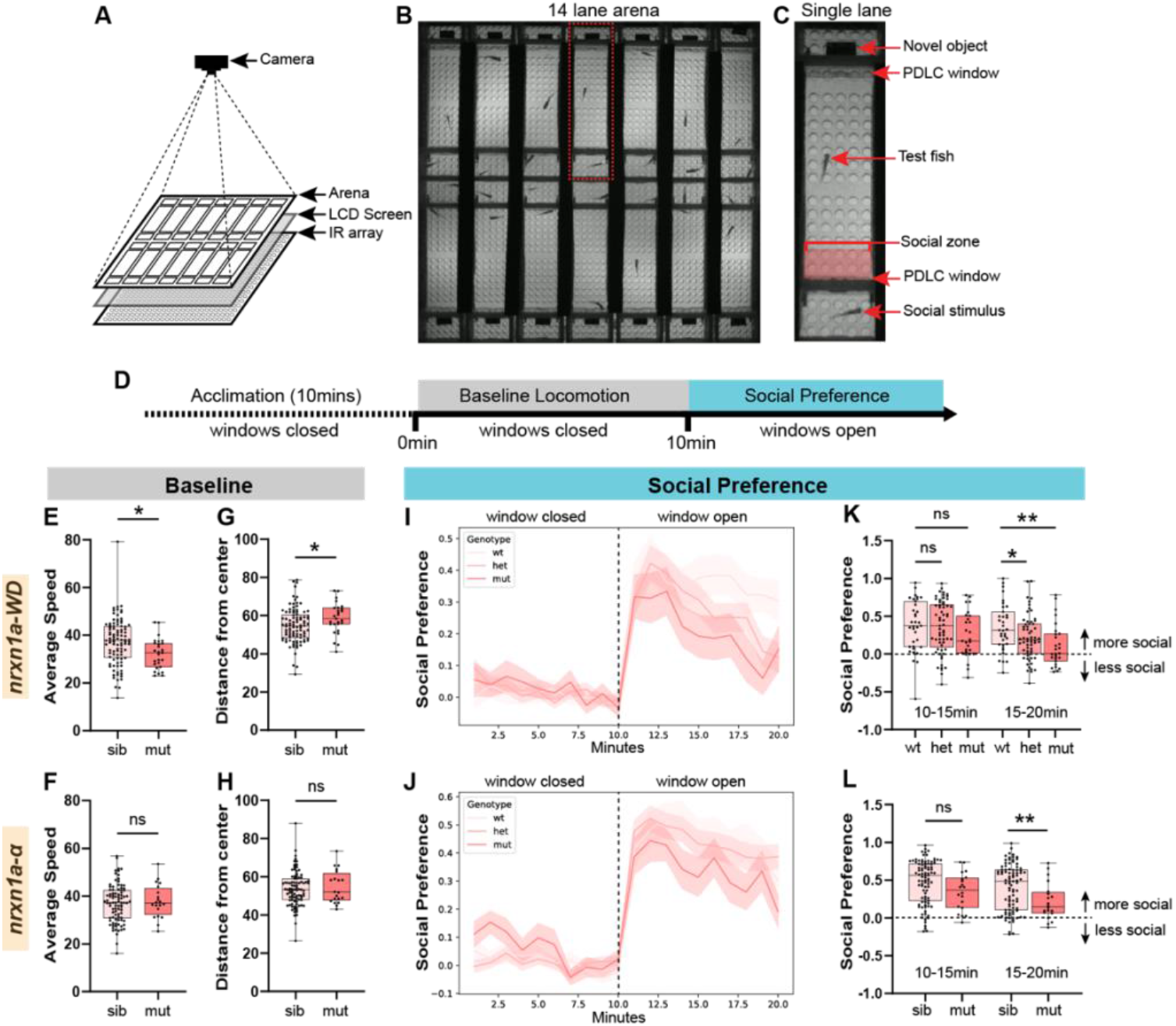
nrxn1a deletions cause social interaction deficits. (A) Juvenile zebrafish behavior apparatus diagram. Arena is illuminated from below by an IR array and an LDC screen. Camera records from above. (B) Still image of the arena from a video recording. (C) Zoom in of one lane of the arena. The test fish resides in the main testing lane which is flanked by two smaller end compartments into which a social stimulus and a novel object are placed. The testing lane is separated by a PDLC window. The social zone (red box) is defined as the area immediately adjacent to the social stimulus. (D) Testing paradigm. Zebrafish are allowed to acclimate to the apparatus for 10mins while the PDLC windows are closed. Baseline locomotion is recorded for 10mins while the PDLC windows remain closed. The PDLC windows open and social preference is assayed for an additional 10mins. (E-H) Average speed (E,F) and average distance from the center of the well (G,H) during the baseline period for *nrxn1a-WD* (E) and *nrxn1a-α* deletion mutants compared to siblings. (I-J) Social Preference over time for *nrxn1a-WD* and *nrxn1a-α* deletion mutants compared to siblings. Note the increase in social preference when the window opens after 10mins. (K,L) Social Preference during the first (10-15min) and second (15-20min) half of the social preference assay for *nrxn1a-WD* and *nrxn1a-α* deletion mutants compared to siblings. *p<0.05, **p<0.01 based on Student’s t-test or One-way ANOVA.

Using this system, we assessed both baseline and social behavior in *nrxn1a-WD* and *nrxn1a-α* deletion lines. Importantly, at 5wpf, homozygous and heterozygous *nrxn1a* animals appeared indistinguishable from age-matched wild-type siblings based on total body length (Fig S3D). To assess if *nrxn1a* function was required for behavior during the baseline period, we measured average speed as well as center-avoidance, as measured by the distance of the fish from the center of the long arm compartment. During the baseline period, neither *nrxn1a-WD* nor *nrxn1a-α* heterozygotes displayed differences from wild-type animals, so these were grouped together as wild-type siblings (Fig S3A-C). Compared to wild-type siblings, *nrxn1a-WD* mutants displayed reduced speed (Fig 6E) and an increase in their distance from the center compared to siblings (Fig 6G). In contrast, we did not observe differences in *nrxn1a-α* mutants for either of these metrics (Fig 6F,H), suggesting that for baseline parameters, other *nrxn1a* isoforms may be able to compensate for *nrxn1a-α* loss. During the social preference period, wild-type sibling animals displayed a robust social preference that slightly decayed over the 10min period (Fig 6I-L). While homozygous *nrxn1a-WD* mutants exhibited a similar initial social preference (Fig 6I,K) as wild-type siblings, it decreased significantly over time and became different from wildtypes in the second half of the assay (Fig 6K). Interestingly, heterozygous *nrxn1a-WD* mutants exhibited an intermediate phenotype that was also significantly different from wildtypes in the second half of the assay (Fig 6K). In contrast, wildtypes and heterozygous *nrxn1a-α* mutants displayed similar social preferences (Fig 6J, S3C) and, though *nrxn1a-α* homozygous mutant animals displayed a significantly reduced social preference compared to siblings (Fig 6L), it was less severe than that of *nrxn1a-WD* mutants. Together, this data defines a clear role for *nrxn1a-α* in social preference but suggests that larger deletions that also encompass the *β* and *γ* isoforms cause more severe phenotypes. Therefore, both our larval and juvenile results indicate that deletions that differentially affect isoforms can have varied effects on behavior, emphasizing the importance of considering deletion type when interpreting clinical alleles.

## Discussion

In this study, we demonstrate that across development deletions of zebrafish NRXN1 isoforms cause both unique and partially overlapping behavioral phenotypes. By generating matched *α-, β-, γ-*, and whole-locus deletions and measuring quantifiable behavioral features using previously validated assays, we show that deletions of isoform-specific exon groups shape discrete behavioral domains, while the full-locus deletion produces combined or more severe phenotypes. These findings provide the first direct comparison of isoform-selective NRXN1 loss *in vivo* and underscore the importance of understanding the specific genomic architecture of NRXN1 deletions when considering pathophysiology, particularly given the substantial inter-individual variability in deletion boundaries.

While mammalian NRXN1 *α, β*, and *γ* isoforms have been previously described, their individual behavioral functions have been incompletely explored. Indeed, most of the *NRXN1* literature to date has focused on *NRXN1*-*α* which has been shown to have important roles in social behavior, locomotion, and anxiety^7–9^. However, there are few reports of specific roles for NRXN1 *β* and *γ*. Recent work in *C. elegans*, which only have *α* and *γ* isoforms, has shown a specific role for *nrx1*-*γ* in a behavioral response to food deprivation^29^, highlighting the need for further studies that address isoform-specific functions. Our study is the first to consider all NRXN1 isoforms in the same experimental paradigm which allows comparison across mutant lines. Similar to prior mouse studies, our findings support a role for NRXN1 in social behavior, locomotion, and thigmotaxis (as measured by distance from the center) which has been previously suggested as a measure of anxiety in zebrafish^30^. Combined, our results support a model in which *α, β*, and *γ* isoforms contribute in both shared and distinct ways which is discussed more thoroughly in the following paragraph. In addition, for each of the *nrxn1a* mutant lines, our data shows that larval behavior (which corresponds approximately to the pre-natal stage in humans^31^) is dysfunctional prior to the emergence of more complex behavioral deficits, supporting a neurodevelopmental origin of *NRXN1* pathogenesis. This contrasts with a recent report indicating that zebrafish lacking both *nrxn1a* and *nrxn1b* do not exhibit detectable behavioral phenotypes at the larval stage^32^. Differences in experimental design, including the temporal resolution of behavioral assays and the use of sibling versus non-sibling controls, may contribute to these differing observations. Notably, our analyses leverage high–temporal resolution behavioral measurements and within-clutch sibling controls, which may increase sensitivity to early developmental phenotypes.

Our larval phenotyping reveals that *nrxn1a-α* and *nrxn1a-β* mutants have opposing behavioral phenotypes in the VMR and DF assays. Additionally, we find that *nrxn1a-WD* mutants, in which both the *nrxn1a-α* and *nrxn1a-β* are deleted, have intermediate VMR and DF phenotypes that are less severe than *nrxn1a-α* mutants. Together, this data supports a model whereby transcripts arising from distinct promoters may drive opposing effects. However, we cannot rule out that isoform-specific deletions may cause changes in splicing or compensatory expression changes of other isoforms which could underlie the observed opposing phenotypes. In contrast to the VMR and DF phenotypes, for the ASR, we only observe a hypersensitivity phenotype for the *nrxn1a-WD* deletion, suggesting *nrxn1a* isoforms may function redundantly to regulate the ASR. Similarly, in the juvenile phenotyping, we observe the same trend, with *nrxn1a-WD* mutants displaying more severe social phenotypes and additional phenotypes that are not present in *nrxn1a-α* mutants. Combined, our data supports a complex picture of *nrxn1a* isoform function, whereby isoforms perform both unique and shared functions that are both behaviorally and temporally specific.

The complexity of our results mirrors the complexity of clinical *NRXN1* deletions. Since *NRXN1* deletions are non-recurrent, non-related individuals have unique deletions. As such, these deletions can have specific effects on NRXN1 function. Previous studies have suggested *NRXN1-α* loss of function caused by deletions in the 5’ region of *NRXN1* as a main mechanism underlying NRXN1’s association with neurodevelopmental phenotypes. However, recent results demonstrate that 5’ and 3’ NRXN1 deletions cause distinct functional outcomes via different mechanisms. Our data supports this view and provides an additional layer of complexity suggesting that the degree to which specific deletions affect NRXN1 isoforms (*α, β, γ*) should be considered.

There are numerous reports of zebrafish social preference assays in the literature, most of which use adult animals, assayed one at a time and usually require physical intervention to either provide or reveal a social cue^25,26^. To solve the need for physical intervention, one group previously employed controllable electrochromic film with good effect^33^. More recently,

moderate-throughput approaches, focusing on juvenile stage animals have been developed^27,28^. However, previous moderate throughput behavioral arenas for social interaction studies in zebrafish have required either netting social stimulus fish into the arena^28^ or physical removal of an opaque separator^27^ for social presentation. To reduce the need for physical interventions that have high potential to interfere with behaviors of interest and to allow for precise control of stimulus presentation, we used Arduino-controlled PDLC films. Compared to electrochromic film, PDLC film is less expensive, responds more instantaneously, and provides improved transparency. Using this approach, we were able to quantify social preference and wild-type animals and observe social phenotypes in *nrxn1a*-deleted animal. By combining PDLC film and a multiplexed setup, this novel platform provides a moderate-throughput system with external, temporal control over stimulus presentation that represents a significant advance in the field.

## Methods

### Experimental model details

Experiments were conducted on 6 dpf larval zebrafish and 5 wpf juvenile zebrafish with body lengths of ∼10-15mm (see Fig S3D) (Danio rerio, TLF strain). Larvae were raised in E3 medium at 29°C on a 14:10 hr light cycle and juvenile and breeding adult zebrafish were maintained at 28°C on a 14:10 hr light cycle. All animal protocols were approved by the University of Pennsylvania Institutional Animal Care and Use Committee (IACUC).

### Crispant experiments

Crispants were generated as described previously^13^. Briefly, three gRNAs targeting three different regions across the *nrxn1a* and *nrxn1b* loci were designed using ChopChop v3 (https://chopchop.cbu.uib.no/) (Supp X). Custom Alt-R CRISPR-Cas9 crRNAs (IDT) were annealed with tracrRNA (IDT, #1072533) to form gRNAs which were subsequently complexed with Cas9 protein (IDT, #1081061) to make the final ribonucleoprotein (RNP) complex. Three non-targeting crRNAs (IDT, #1072544, 1072545, 1072546) were used to make the RNP for controls. Single-cell wildtype (TLF) zebrafish embryos were then microinjected within 15minutes of fertilization with 1nl of RNP mix containing 357pg (10.1fmol) of each gRNA and 5029pg (30.5fmol) of Cas9. Embryos displaying acute toxicity or damage from microinjection were removed from analysis. Behavior of crispants was then assayed at 6dpf as described below.

### Stable deletion line generation and genotyping

Mutant alleles were generated using CRISPR-Cas9 mutagenesis. Two gRNAs flanking the desired region, along with a third nested gRNA within the region to be deleted, were designed using ChopChop v2 (https://chopchop.cbu.uib.no/). RNP complexes were then made and injected into single-cell wildtype (TLF) zebrafish embryos as described in the above crispant section. F_0_ injected larvae were raised and outcrossed to identify and establish heterozygous carrier lines. Mutant lines were identified and subsequently genotyped by PCR, using primers flanking the outermost gRNA target sites. Allele sequences were obtained by Sanger-sequencing of resulting PCR products. For each allele, genotyping was performed using the two primers flanking the deletion to amplify the mutant allele together with a third nested primer to amplify the wildtype allele (Supp X). Heterozygous deletion carriers were in-crossed to generate wildtypes, heterozygotes, and homozygotes for larval and juvenile behavioral experiments. All behavioral experiments were performed on F_3_ generation animals or later.

### HCR fluorescent in situ hybridization

Endogenous *nrxn1a-α* and *nrxn1a-β* transcripts were detected using HCR fluorescent in situ hybridization (Molecular Instruments, Los Angeles, CA, USA). Probes for *nrxn1a-α* were designed against the *α*-specific exons 1-4. Probes for *nrxn1a-β* were designed against the lone *β*-specific exon. HCR buffers, probes, and hairpins were purchased from Molecular Instruments. Wild-type TLF larvae at 6dpf were fixed with 4% paraformaldehyde in Dulbecco’s phosphate-buffered saline overnight at 4°C and staining was performed as previously described^34^.

### Larval behavioral experiment and analysis

Crispants and stable mutant lines were assayed for sensorimotor phenotypes as previously described^14,15^. Briefly, larvae were arrayed in a 100-well plate and assayed for the VMR, Light flash response, Dark flash response, and acoustic startle response. Recorded videos were then tracked and analyzed offline using previously published Python codes^14^.

### Juvenile behavior arena

The testing arena consists of a 2×7 array of testing lanes and is made of laser-cut acrylic. Each testing lane is 80mm X 30mm (holds ∼18ml of water) and is flanked on each end by a compartment of 18mm X 30mm (holds ∼5ml of water). The bottom of the arena is clear transparent acrylic which allows fish to observe an LCD screen that is placed underneath the arena and backlit by LED white lights. An IR array also illuminates from below which is detected by a Chameleon3 monochrome camera mounted above (CM3-U3-13Y3M-CS, Teledyne FLIR) and fitted with an IR longpass filter (LP800-30.5, Midwest Optical) at 20fps. Side walls of the arena are white opaque acrylic to prevent fish in adjacent lanes from visualizing one another. The end walls of the lanes are fitted with PDLC windows (MEGICOLIM) which are opaque but become transparent when triggered by an Arduino, allowing fish to observe what is placed in the end compartments (i.e. social stimulus fish or novel object). The entire testing arena is contained in a black-out curtain enclosure to minimize external interference. Information regarding hardware and code for the juvenile behavioral arena is openly accessible at https://github.com/pdcampbell/ZF-juvenile-rig.

### Juvenile behavior experiment and analysis

Heterozygous deletion carriers were in-crossed and progeny were raised to 5wpf alongside age-matched wild-type TLF fish to serve as social stimuli. 5wpf fish were moved from their housing tanks and placed into either the testing lanes (deletion fish to be tested) or the end compartments (wild-type social stimuli). Fish were allowed to acclimate to the environment for 10 mins while white light was projected from below by the LCD screen. Then, baseline movements were recorded for 10mins. PDLC films were then triggered to open and social preference was recorded for an additional 10mins. Tested fish were then individually removed from the testing arena and genotyped. Offline tracking of recorded videos was performed with custom written Python codes. To identify objects in each well, each image of the video was background subtracted, Gaussian blurred and thresholded. Fish that were improperly tracked (i.e. no objects were detected in the well or more than one object was detected) in >10% of the frames or that have >200 consecutive improperly tracked frames were discarded (∼5% of fish). Remaining fish displayed a high tracking accuracy (>99% of frames tracked correctly, over n=254 fish). Missing frames were filled in using the scipy.interpolate function. Fish that did not move for >1000 consecutive frames (i.e. 50s) during the baseline period were also excluded from analysis as they were deemed to be exhibiting freezing behavior. Excluded fish spanned all genotypes and were not clustered in one group. Object centroids for remaining fish were then used to calculate of average speed, position in the well relative to the social stimulus, and position in the well relative to the center. Average speed was computed as the distance travelled by the centroid over the time spent moving. Social preference (SP) was calculated for each fish by subtracting the fraction of time in the area nearest the social stimulus (social zone) during the last 5mins of the baseline period when the PDLC is closed from the fraction of time in the social zone when the PDLC is open. Distance from the center was computed as the distance between the centroid of the fish and the center of the well. Data was binned into 1minute bins with an average computed for each 1min bin. Fish lengths were measured from the videos in ImageJ. Information regarding tracking and analysis code for the juvenile behavioral experiments is openly accessible at https://github.com/pdcampbell/ZF-juvenile-rig.

## Acknowledgements

The authors would like to acknowledge the University of Pennsylvania Penn Electronic Design Shop Core Facility (RRID:SCR_021107), the University of Pennsylvania Cell and Developmental Biology Microscopy Core, and the Penn Zebrafish Facility. This work was supported by grants to P.D.C. (NIH K08NS135125), M.G. (NIH R01NS118921), and the University of Pennsylvania Autism Spectrum Program of Excellence.

---

**Fig S1.**
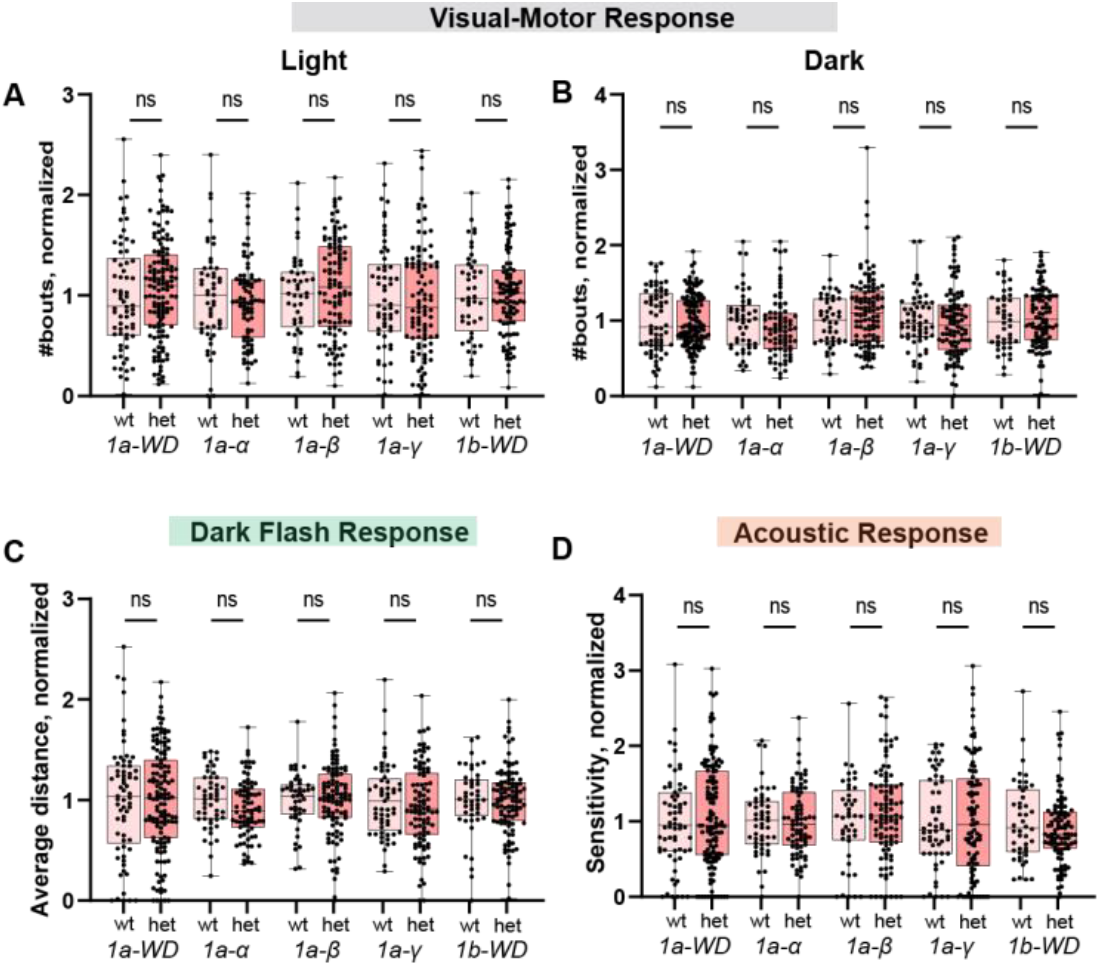
(A,B) Average of normalized bouts across all light and dark minutes for each deletion line, heterozygous mutants compared to wildtypes. (C) Average distance travelled following a flash of darkness, normalized to wildtypes for each deletion line, heterozygous mutants compared to wildtypes. (D) Sensitivity to acoustic stimuli, normalized to wildtypes for each deletion line, heterozygous mutants compared to wildtypes. Numbers (n) can be found in legend of Fig 5. Statistics based on Student’s t-test.

**Fig S2.**
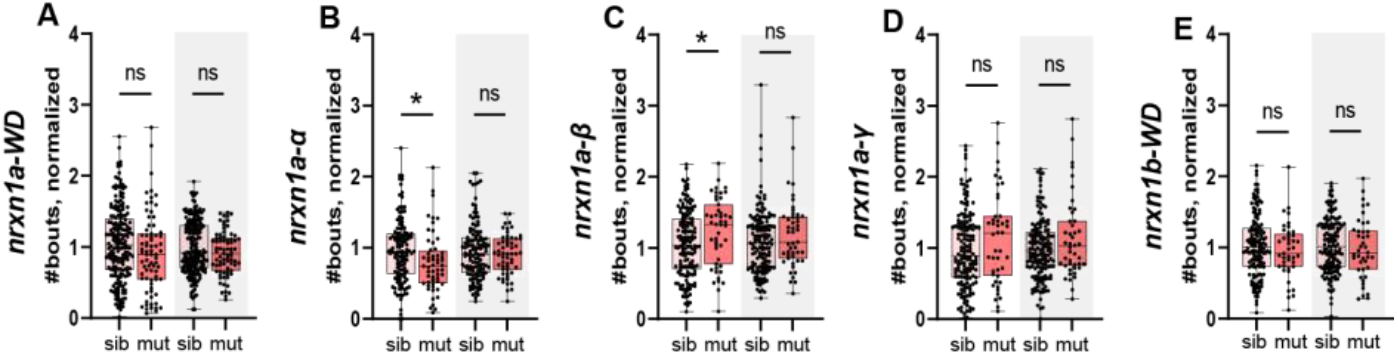
(A-E) Average of normalized bouts across all light (white) and dark (gray) minutes for each deletion line. Numbers (n) can be found in legend of Fig 5. Statistics based on Student’s t-test. *p<0.05 based on Student’s t-test.

**Fig S3.**
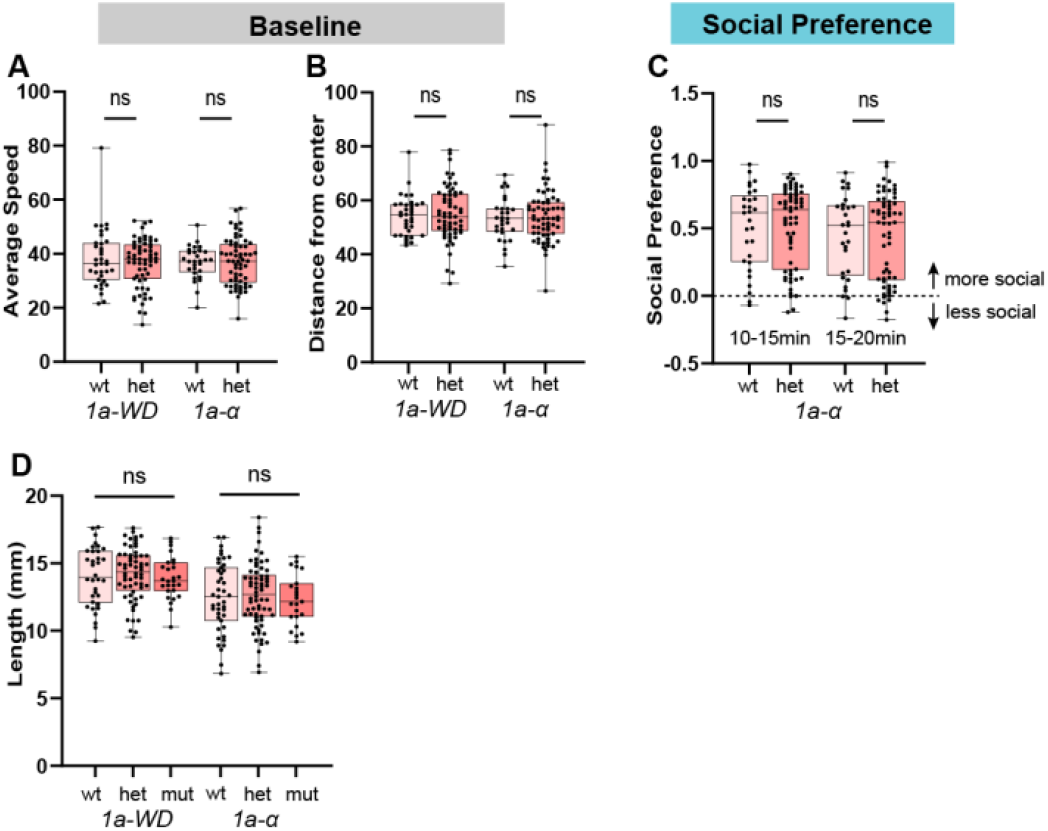
(A,B) Average speed (A) and average distance from the center of the well (B) during the baseline period for *nrxn1a-WD* (E) and *nrxn1a-α* deletion heterozygous mutants compared to wildtypes. (C) Social Preference during the first (10-15min) and second (15-20min) half of the social preference assay for *nrxn1a-WD* and *nrxn1a-α* deletion heterozygous mutants compared to wildtypes. (D) Length of zebrafish for *nrxn1a-WD* and *nrxn1a-α* deletion mutants compared to wild-type and heterozygous siblings. Stats based on Student’s t-test or One-way ANOVA.

